# FINDEL: A Deep Learning Approach to Efficient Artifact Removal From Cancer Genomes

**DOI:** 10.1101/2022.11.12.516244

**Authors:** Denis Tan, Pengfei Zhou, Shaoting Zhang, VicPearly Wong, Jie Zhang, Edwin Long

**Affiliations:** New Silkroutes Group Ltd, 119962, Singapore; SenseTime Research, Shanghai 200233, China; Qing Yuan Research Institute, Shanghai Jiao Tong University, Shanghai 200240, China

**Keywords:** bioinformatics software, cancer genomics, deep learning, machine learning, mutational signatures, sequencing artifacts, somatic mutations, variant refinement

## Abstract

Next-generation sequencing technologies have increased sequencing throughput by 100-1000 folds and subsequently reduced the cost of sequencing a human genome to approximately US$1,000. However, the existence of sequencing artifacts can cause erroneous identification of variants and adversely impact the downstream analyses. Currently, the manual inspection of variants for additional refinement is still necessary for high-quality variant calls. The inspection is usually done on large binary alignment map (BAM) files which consume a huge amount of labor and time. It also suffers from a lack of standardization and reproducibility. Here we show that the use of mutational signatures coupled with deep learning can replace the current standards in the bioinformatics workflow. This software, called FINDEL, can efficiently remove sequencing artifacts from cancer samples. It queries the variant call format file which is much more compact than BAM files. The software automates the variant refinement process and produces high-quality variant calls.

## 1 Introduction

Since the discovery of the DNA structure in 1953 [1], significant progress in genetic understanding has been made over the past few decades. The advent of DNA sequencing instruments made possible the Human Genome Project (HGP) which was a global research initiative in 1990 to sequence the entire human genome consisting of 3 billion nucleotide base pairs [2, 3]. The international and cross-disciplinary nature of HGP has popularized the idea of open-source software and data accessibility where scientists and researchers all over the world have equal access to the ever-increasing repository of human genomes. The availability of individual genome data through cloud infrastructure and technologies has revolutionized the field of precision medicine where scientists can mine valuable information and recommend personalized treatment [4]. This can greatly improve the healthcare standards of the current and future generations.

The completion of the HGP has led to higher demand for sequencing technologies and datasets to solve more sophisticated biological problems. However, those efforts were hampered by the relatively low throughput and high costs of sequencing at that time [5]. This was changed in the mid-2000s with the invention of the high-throughput sequencing platform which resulted in a 50,000 times decrease in the cost of sequencing the entire human genome [6]. The promising results led to the term “next-generation sequencing” (NGS) which refers to platforms capable of massively parallel sequencing technologies that enable ultra-high throughput, scalability, and speed [7]. NGS technologies have increased sequencing throughput by 100-1000 folds [8] and subsequently reduced the cost of sequencing a human genome to approximately US$1,000. These advances have made possible the use of sequencing technologies in clinical settings [6]. NGS technologies have found large-scale usage in de novo sequencing [9], mapping of diseases [10], quantifying expression levels using RNA sequencing [11, 12, 13], and conducting population genetic studies [14, 15, 16].

NGS technologies generate millions to billions of short-read sequences with read lengths of around 75-300 base pairs (bp). More advanced NGS technologies (PacBio, Nanopore, 10x Genomics) are capable of much longer read sequences of more than 10 kilobases [17]. NGS technologies, while cheap and fast, are not flawless. The sequencing procedure is just the initial step of a typical bioinformatics pipeline that spans multiple phases. The raw sequence reads produced by the sequencing machine is stored in a FASTQ or unaligned Binary Alignment Map (uBAM) format. These formats are text-based which store information on the sequence reads such as read identifiers and base quality scores. Next, the reads are aligned to a reference genome and the related metadata are stored in either a Sequence Alignment Mapping (SAM), Binary Alignment Mapping (BAM), or CRAM file formats [18]. These files contain alignment characteristics such as matches, mismatches, and gaps represented in the Concise Idiosyncratic Gapped Alignment Report (CIGAR) format [19]. The BAM or equivalent files are consumed downstream through variant calling algorithms to identify genetic mutations such as single nucleotide polymorphisms (SNPs) consisting of transversions and transitions, insertions and deletions (InDels), and tumor mutation burden [20, 21]. The identified variants are stored in a tab-delimited text file format commonly known as the Variant Call Format (VCF).

Due to the downstream implications of NGS data, the inaccurate reads can cause erroneous identification of variants, specifically false positives (FP) and false negatives (FN). NGS data are susceptible to errors due to a myriad of factors such as base-calling and alignment errors [22]. This is further exacerbated by the lack of standardization in bioinformatics pipelines which are crucial to obtaining the correct data interpretation and clinical insights [23]. Errors in variant calling can result in missed detection of mutations which can be disastrous. Mutations are the cause of several diseases especially cancer [23].

A major shortfall of NGS technology is the frequent incorrect scoring of bases attributed to the existence of artifacts during the sample preparation and sequencing stage [24]. Heterogeneous mixtures can suffer from amplification bias during the polymerase chain reaction (PCR) process which produces skewed populations [25]. Other types of polymerase mistakes during the PCR stage such as base misincorporations and rearrangements because of template switching can also cause inaccurate variant calls. Cluster amplification, sequencing cycles, and image analysis are also prone to mistakes contributing to roughly 0.1-1% of bases being erroneously called [26]. These artifacts present a challenge for calling rare genetic variants as deep sequencing is ineffective when the base call error rate is high. This effectively limits the application of NGS in fields such as metagenomics [27, 28], forensics [27], and human genetics [29, 30]. The accuracy demands of some clinical applications are even higher where the base calling error rate has to fall below 1 in 10,000. Examples of these clinical applications include detection of circulating tumor DNA [31], monitoring of response to chemotherapy using personalized tumor biomarkers [32], and prenatal screening for fetal aneuploidy [33, 34]. Due to these artifacts, the common NGS technologies suffer from a base-calling error rate of around 1 in 100 which severely falls short of the standard required by these clinical applications.

Automated pipelines to facilitate reads-to-variants workflow are widely available in this day and age, a prominent one being the Genome Analysis Toolkit (GATK) Best Practices Workflows which contain step-by-step recommendations for conducting variant discovery analysis using NGS data. These pipelines have built-in filters to remove false variant calls from sequencing errors, read misalignments, and other types of errors. However, there is little support for the efficient removal of artifacts due to variant caller inaccuracies [35]. According to the Association for Molecular Pathology (AMP) guidelines for interpretation and annotation of somatic variation, it is crucial to perform additional refinement of somatic variants to remove variant caller inaccuracies as these can result in suboptimal patient management and therapeutic opportunities [36, 37]. As of now, the standard practice for somatic variant refinement is to manually inspect the variants and this is usually performed by trained personnel such as bioinformaticians. Manual inspection is useful for incorporating domain knowledge commonly excluded by automated variant callers such as Mutect [38], SomaticSniper [39], Strelka [40], and VarScan2 [41]. They can detect inaccurate variant calls due to amplification bias of small fragments, errors in sequencing reads, and poor alignment in certain areas [35]. Efforts have been made to develop automated methods for variant refinement but progress is slow due to computational limitations. Manual inspection of variants for additional refinement is still necessary for high-quality variant calls and subsequent downstream analyses [42].

Although manual inspection of variants has been utilized for several years in clinical diagnostic and molecular pathology applications [43, 44, 45], somatic variant refinement strategies are not documented extensively and reported minimally by studies involving postprocessing of automated variant calls [45, 46, 47, 48, 49]. These results in a lack of standardized protocol for somatic variant refinement, increased variability of work performed by different labs, and difficulty in reproducing the work of others [49]. Additionally, manual inspections are usually conducted on BAM files using genomic viewers such as Integrative Genomics Viewer (IGV) [50, 51], Savant [52], Trackster [53], and BamView [54]. These BAM files are extremely large and require sophisticated storage solutions. This is because the format stores data on a per-read basis and the space requirement grows almost linearly with the number of reads [55]. A BAM file for a 30x whole genome requires about 80-90 gigabytes of storage. A lab or research institution which handles around 1000 samples needs to secure approximately 80 terabytes of disk space [56]. The sheer size of these BAM files impedes the progress of the manual inspection process which will require more labor and time. A more efficient and scalable variant refinement solution with light processing and storage requirements while eliminating variability in the downstream analyses needs to be developed.

A promising area of research beneficial to variant refinement lies in the study of mutational signatures. Mutational signatures have been linked to mutational processes which drive somatic mutations in cancer genomes [57]. The Pan-Cancer Analysis of Whole Genomes (PCAWG) Consortium [58] of the International Cancer Genome Consortium (ICGC) and The Cancer Genome Atlas (TCGA) analyzed 84,729,690 somatic mutations identified from 4,645 whole-genome and 19,184 exome sequences that cover a wide range of cancer types and generated many mutational signatures [59]. The mutational signatures consist of 49 single-base-substitution (SBS), 11 doublet-base-substitution (DBS), 4 clustered-base-substitution, and 17 small insertion-and-deletion (Indel) signatures. Further classification of each mutation type was performed whereby SBSs consisted of 96 classes, DBSs consisted of 78 classes, and Indels consisted of 83 classes. These mutational signatures can be found in the Catalogue of Somatic Mutations in Cancer (COSMIC) database [60]. The PCAWG study also discovered that DNA sequencing artifacts, analysis artifacts, technical artifacts, variability in NGS technologies, and usage of different variant calling methods can generate characteristic mutational signatures as well [59]. These discoveries open up the possibility of using mutational signatures to filter artifact-mediated variant calls from true variants.

Our team proposes FINDEL (Find & Delete), a variant refinement software written in Python programming language that removes sequencing artifacts from human cancer samples using mutational signatures and deep learning. Along with each sample, FINDEL generates a Hypertext Markup Language (HTML) report and 2 separate VCF files containing refined and artifactual mutations respectively. FINDEL is light on computational requirements and is capable of running on a simple local machine. The performance and speed of FINDEL are validated using open-source datasets.

## 2 Methods

### 2.1 Overview of FINDEL

The main function of FINDEL is to automate the process of removing sequencing artifacts using mutational signatures from the COSMIC database coupled with a deep learning approach. The software is intended for use in the variant refinement stage and is not meant to replace the variant calling algorithms. FINDEL can remove variability in the bioinformatics pipeline within the post variant calling phase. Currently, FINDEL focuses on artifacts involving single-base substitutions (SBSs) or also known as single-nucleotide polymorphisms (SNPs). SNPs form the majority of the genetic variations found in the human genome. In a typical human genome, more than 99.9% of variants are characterized as SNPs and short indels with SNPs being 6-7x higher in frequency than short indels [61]. SNPs are also responsible for the onset of multiple cancer types including breast cancer [62], chronic lymphocytic leukemia [63, 64, 65, 66, 67, 68], neuroblastoma [69], gastric carcinogenesis [70, 71, 72, 73, 74, 75, 76], prostate cancer [77], and others [78, 79].

### 2.2 Identification of Single-Base Substitutions Mutational Signatures Linked to Sequencing Artifacts

The initial step in building the artifact removal algorithm is to identify the SBS mutational signatures that are linked to the sequencing artifacts. These can be found in the COSMIC database. A total of 18 SBS mutational signatures have been classified as possible sequencing artifacts: SBS27, SBS43, and SBS45-SBS60. The remaining signatures either have valid proposed aetiologies or unknown causes. As of now, SBS signatures can be segregated into 96 different contexts. There are 6 possible base substitutions: C>A, C>G, C>T, T>A, T>C, and T>G. Each base substitution can be further analyzed in its 5’ and 3’ nucleotide context forming a total of 96 trinucleotide contexts [80].

### 2.3 Input Requirements for Algorithm

FINDEL takes in a VCF file containing a single sample. PyVCF module in Python is used to parse the VCF file for further processing steps. The other input required by the algorithm is the reference sequence. The reference sequence is used to extract the mutation type and reference context.

### 2.4 Output Files

FINDEL outputs 2 separate VCF files containing the refined and artifactual mutations respectively. The refined VCF file can be used for downstream analyses.

### 2.5 Supervised Deep Learning

After the information from the mutational signatures are obtained, we propose a supervised deep neural network-based method to directly predict the mutation types from the VCF file. We devised the specific modules to deal with a variety of features in addition to the mutational signatures and utilized a complex network to fuse the diverse information for the mutation site identification. The deep learning architecture has a comprehensive embedding representation to enhance the algorithm performance. Upon training completion, FINDEL can conduct inference efficiently without the need for a customized variant refinement optimization process for every new cancer sample.

#### 2.5.1 Data Preprocessing

Firstly, we constructed the features of mutation types. All point mutations are classified into one of the 96 SBS nucleotide contexts, represented by an index ranging from 0 to 95. Secondly, we parsed the site features. Specifically, we used the “INFO” and “SAMPLE” columns from the provided input VCF file. One-hot encoding and standard scaling were used to preprocess the discrete and continuous features respectively. Thirdly, we built context features. We selected 10 bases at the left and right of each mutation site (a total of 21 bases). Each base is represented by a number (T:0, C:1, A:2, G:3, others:4), and the context feature was represented as a 21-dimensional vector. Finally, we adopted the Hap.py tool to process the original annotation file as the ground-truth labels in this study.

#### 2.5.2 Neural Network Architecture

Fig. 1 shows the proposed neural network architecture. We used a learnable embedding layer to map the 96-dimensional mutation type into a high-dimensional space. Each mutation type is converted into a point in the high-dimensional space, the coordinates of which constitute an embedding vector for the type. The parsed site features were combined and fed into a linear layer. The reference context is passed into another embedding layer, converting the 21 bases in the mutation neighborhood into a two-dimensional context matrix. We used a two-layer 1D-convolution to model the multiple sequences and extract the backward and forward relationships in the context matrix. The context matrix was transformed into a vector.

**Fig. 1.**
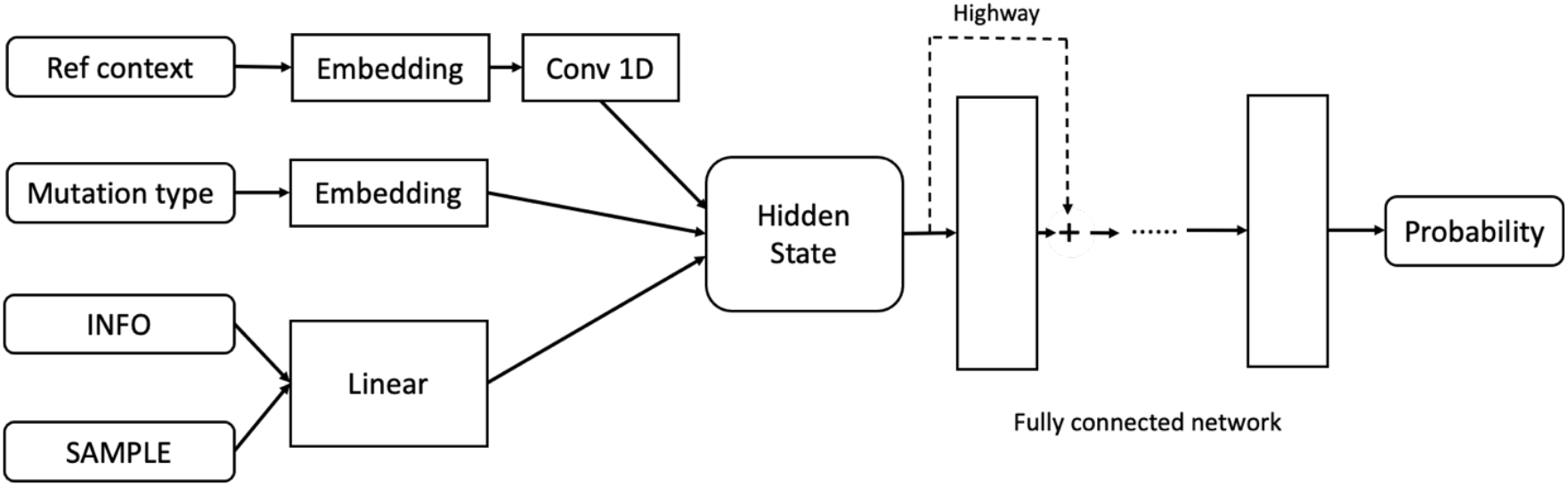
Neural network architecture.

**Fig. 2.**
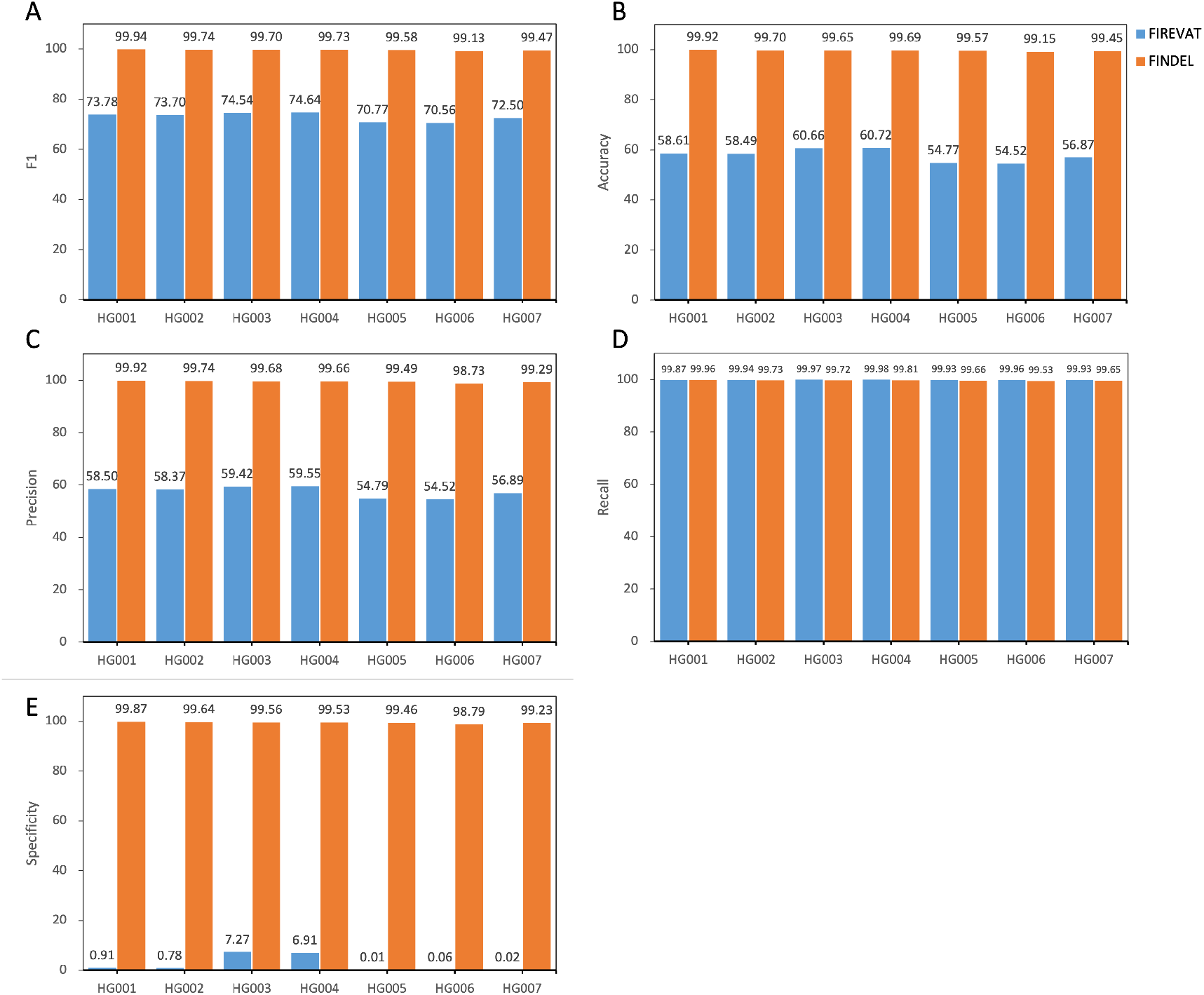
Performance comparison between py_artifact_removal (before deep learning) and DL_v11 (after deep learning) based on 5 metrics. All values are in percentages unless otherwise indicated. All datasets used here are from the GIAB consortium reference samples (HG001-HG007). The PY_ARTIFACT_REMOVAL method is based on mutation signatures, which tends to produce positive results. The supervised learning-based DL-V11 method is able to accurately identify artificial mutations. In all four metrics except recall, dl-v11 has a huge improvement.

**Fig. 3.**
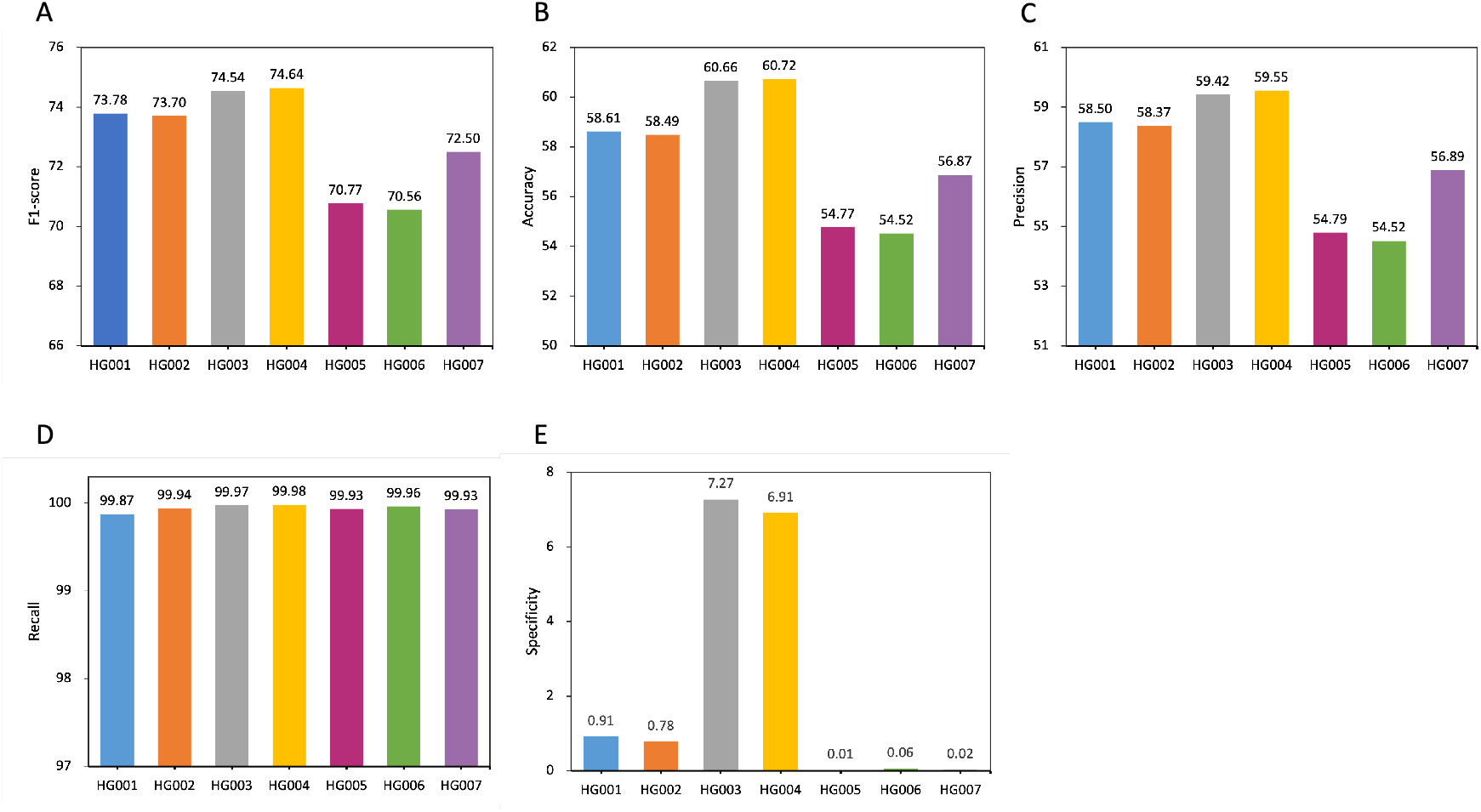
Performance comparison within py_artifact_removal (before deep learning) based on 5 metrics. All values are in percentages unless otherwise indicated. All datasets used here are from the GIAB consortium reference samples (HG001-HG007). Different races will affect the performance of the model. PY_ARTIFACT_REMOVAL has better performance in Utah/European (HG001) and Ashkenazi Jewish (HG002/HG003/HG004) than Han Chinese (HG005/HG006/HG007) ancestries.

**Fig. 4.**
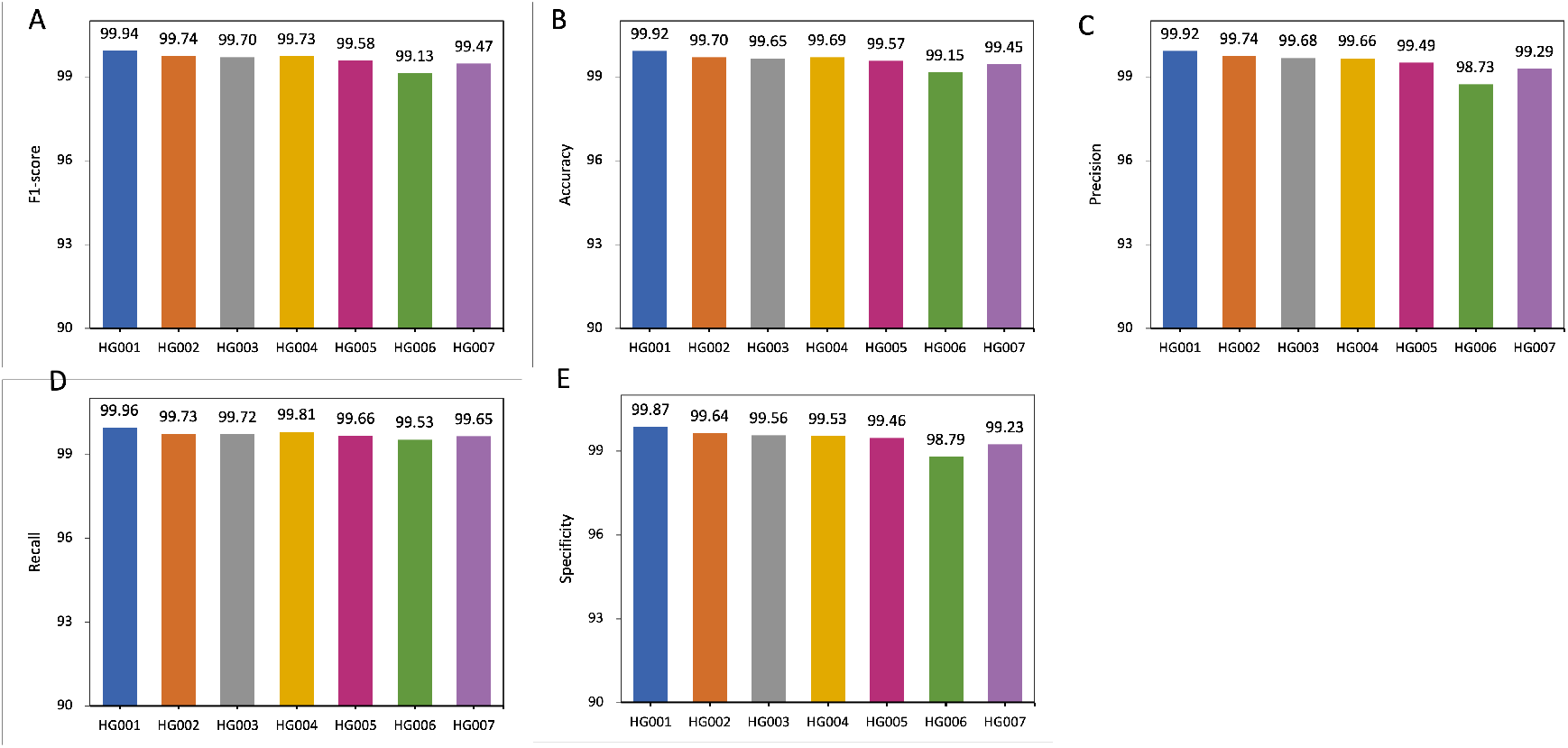
Performance comparison within DL_v11 (after deep learning) based on 5 metrics. All values are in percentages unless otherwise indicated. All datasets used here are from the GIAB consortium reference samples (HG001-HG007). An independent model is trained for each sample. DL_v11 achieves the best performance on HG001, with an f1-score of 99.94. Lower scores were achieved on the other human genomes with the Han Chinese(HG006) ancestry genome performing the poorest.

**Fig. 5.**
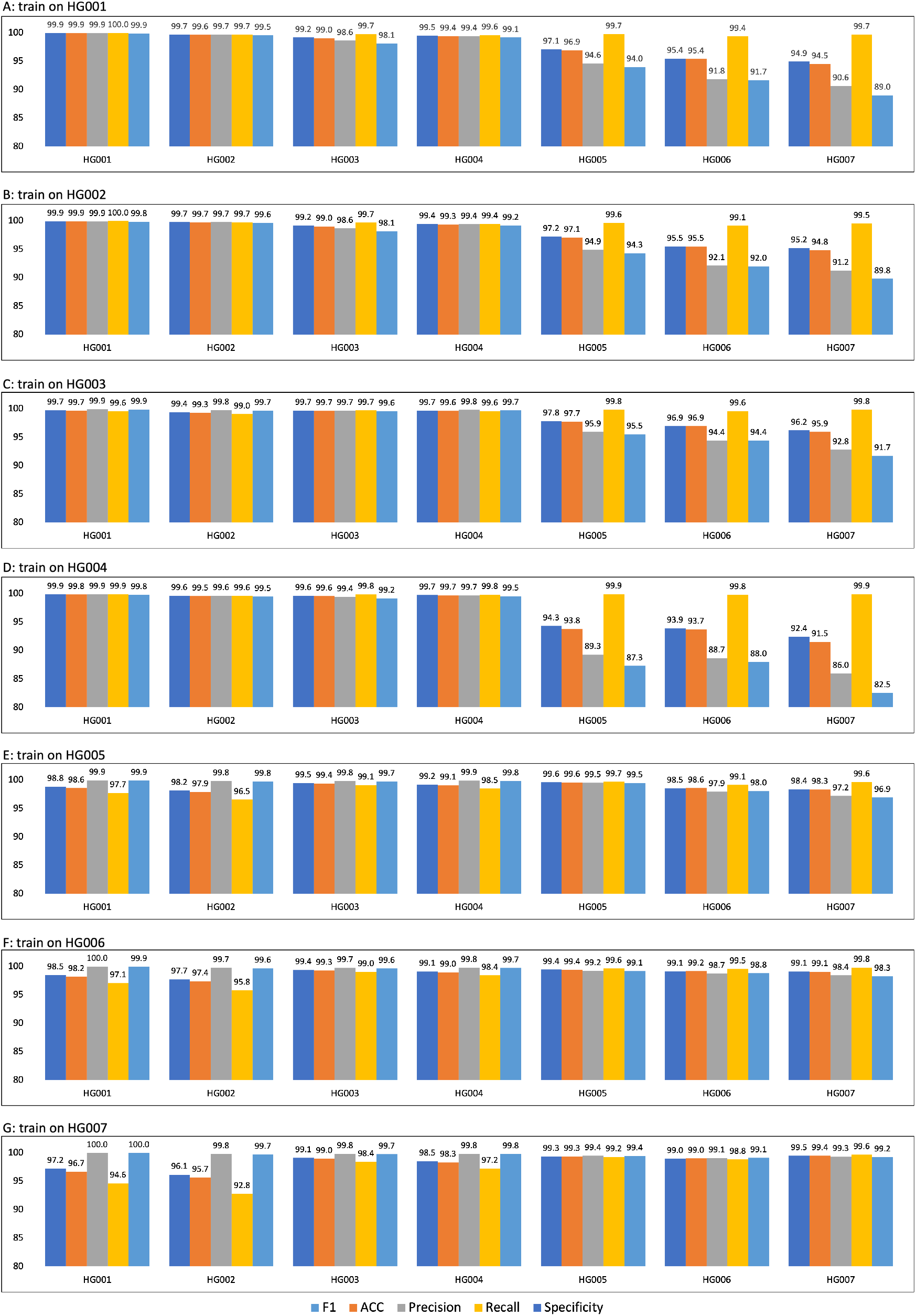
Performance comparison within DL_v11 (after deep learning) for different trainsets. The model achieves better performance when the test and training data come from the same sample. When the training and test data come from different sources, the algorithm performance will be affected. For example, in A Barplot, the training data is chromosome 1 to 19 of HG001(Utah/European), and the test data is chromosome 20 of HG001 to HG007. On HG001 all five metrics achieve a performance of 99.9, but the model predicts more false-positive mutations on HG005/HG006/HG007 (Han Chinese) of different ethnicities, resulting in a lower Specificity

**Fig. 6.**
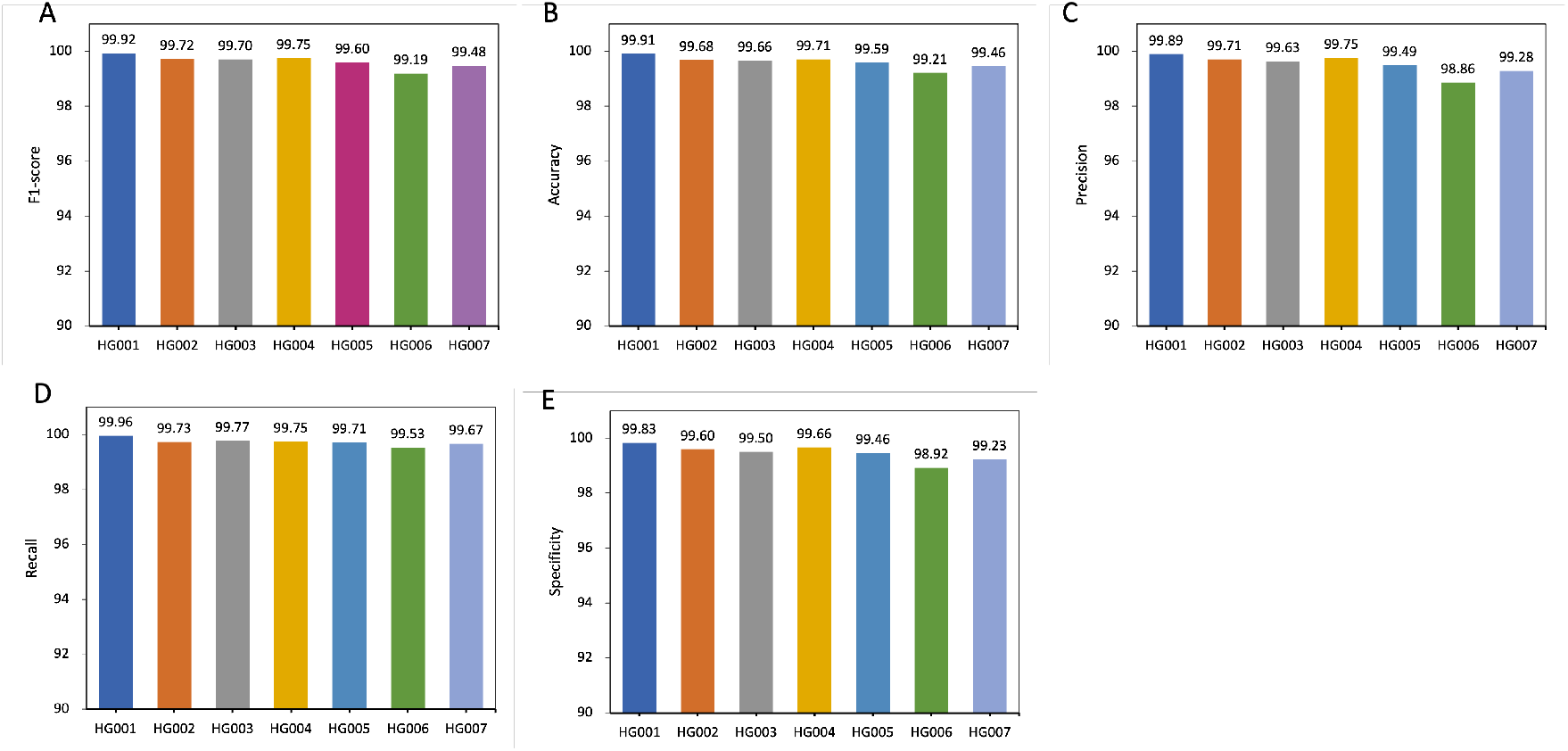
Performance comparison within DL_v11 (after deep learning) for whole trainsets. Training using data from multiple samples was able to improve the robustness of the model and alleviate the effects of different sources of data on model performance. The DL_v11 model trained using chromosome 1 to 19 from HG001 to HG007, the model was able to achieve an f1-score of 99% on all samples.

After extracting the mutation types, site features, and context features, we concatenated the embedding vectors and fed them into a five-layer fully connected network. Each layer consisted of dropout linear transformation, ELU (Exponential Linear Unit) activation function, and highway module. The dropout was used at the beginning of each layer to randomly set the unit weights to 0, helping the model avoid overfitting specific features. Equations (1), Equations (2), and Equations (3) show the formulas for the linear transformation, activation function, and highway module respectively:

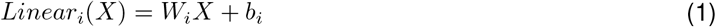

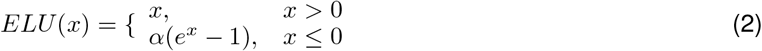

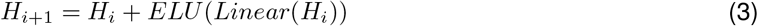

where W and b are learnable parameters. The ELU hyperparameter *α* controls the value at which the activation function saturates for negative inputs. The highway module connects the input of each layer to the end, solving the gradient vanishing problem caused by overly deep networks.

Equation (4) defines the cost function.

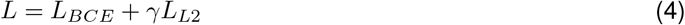

*L_BCE_* refers to the binary cross-entropy loss while *γL*_*L*2_ is the L2 regularization which adds a penalty term to reduce overfitting. We adopted the mini-batch gradient descent algorithm for training and each batch consisted of 5000 variants. The Adam optimizer was used to update the network weights. All hyperparameters were tuned using grid search and evaluated on the validation set. The GIAB consortium reference samples (HG001-HG007) were used for training and testing. Two different training methods were used. The first method built 1 model using 1 VCF file. Within each file, the SNPs were split such that 60% of them were part of the train set, 20% of them were part of the validation set, and 20% of them were part of the test set. The second method used all the SNPs from 1 VCF file and tested them on all the remaining files. For example, a model was trained on all the SNPs from the HG001 VCF file and evaluated on the SNPs from the HG002-HG007 files. Table I shows the final values chosen for the hyperparameters.

### 2.6 Evaluation Data Sources and Processing

The performance and speed of FINDEL are evaluated using high-quality open-source datasets. For the proof of concept training and validation, we will be using 4 different datasets containing a total of 33 VCF files.

#### 2.6.1 HCC1954 Cell Line

This breast cancer cell line whole genome sequencing (WGS) data from a 61 years old Asian female was initiated on October 30 1995 and took around 4 months to establish [81]. We downloaded the VCF file and the benchmarking results from the International Cancer Genome Consortium Data Portal [82]. The reads were aligned to the hs37d5 reference genome. The benchmarking results containing the true positive and false positive variants were stored in 2 mutation annotation format (MAF) files which are tab-delimited text files with aggregated mutation information on a project level. No processing was required since the VCF file was provided directly. A total of 1 VCF file was obtained.

#### 2.6.2 Formalin-Fixed Paraffin-Embedded (FFPE) and Fresh Frozen Whole Exome Sequencing (WES) Samples

This study was part of an attempt by Samsung Medical Center to compare the WES data between FFPE and fresh frozen samples. The datasets were obtained from the Sequence Read Archive (SRA) [83] with the accession number PRJNA301548. The samples were sequenced using the Illumina HiSeq 2000 sequencing system. We used the DNA sequencing data from the following 3 runs: SRR2911437, SRR2911438, and SRR2911453. The paired-end reads were aligned to the HG19 reference genome using the Burrows-Wheeler Alignment tool [84] with the Maximal Exact Match option, more commonly know as the BWA-MEM algorithm. After the reads were aligned, Mutect2 was chosen for the variant calling process. The tumor-only mode with default settings was used. A total of 3 VCF files was obtained.

#### 2.6.3 Whole Exome Sequencing of Biological and Technical Replicates in Breast Cancer Samples

This study was conducted by the Memorial Sloan Kettering Cancer Center on 3 Feb 2016. The datasets were obtained from the SRA with the accession number SRP070662. The samples were sequenced using the Ion PGM sequencing system We used the DNA sequencing data from the following 22 runs: SRR3182418, SRR3182419, SRR3182421, SRR3182423, SRR3182424, SRR3182427, SRR3182429, SRR3182430, SRR3182431, SRR3182432, SRR3182436, SRR3182437, SRR3182440, SRR3182442, SRR3182444, SRR3182446, SRR3182447, SRR3182474, SRR3182475, SRR3182476, SRR3182477, and SRR3182478. The processing steps are the same as the FFPE and fresh frozen WES datasets. A total of 22 VCF files was obtained.

#### 2.6.4 Genome in a Bottle (GIAB) Consortium Reference Samples

The GIAB consortium is jointly hosted by the National Institute of Standards and Technology (NIST) and the Joint Initiative for Metrology in Biology (JIMB) to develop the technical infrastructure and improve clinical insights from whole human genome sequencing. GIAB has a total of 7 human genomes: 1 pilot genome of Utah/European ancestry (NA12878/HG001) from the HapMap project, and 2 son/father/mother trios of Ashkenazi Jewish (HG002/HG003/HG004) and Han Chinese (HG005/HG006/HG007) ancestry from the Personal Genome Project. Each human genome is also accompanied by benchmark variant calls and regions to validate variant calling pipelines. These are considered the gold standard benchmarks within the bioinformatics community. The processing steps are mostly the same as the FFPE and fresh frozen WES datasets with some differences. The b37 reference genome was used in place of the HG19 reference genome. Both originate from the GRCh37 reference genome with some minor differences. However, from a bioinformatics pipeline technical perspective, these variations of reference genomes are not directly interchangeable due to contig name changes. The discrepancies are further elaborated in. The change was due to the addition of the germline resource and the panel of normals (PoN) during the Mutect2 variant calling process. Both VCF files used the b37 reference genome. Therefore, the alignment phase required the use of the same reference genome as well. A total of 7 VCF files was obtained.

The HCC1954 cell line, FFPE and fresh frozen WES, and WES of biological and technical replicates in breast cancer datasets lack the labeling of true mutations. DL_v11 used an unsupervised training method with a negative objective function as the loss function. In the GIAB annotated dataset, we used chromosomes 1 to 19 as the training set, chromosomes 21 and 22 as the validation set, and chromosome 20 as the test set. We used cross-entropy as the loss function,

## 3 Results

FINDEL will be evaluated based on its performance and speed on the evaluation data. For performance evaluation, benchmark VCF files are required as they contain the ground truth variant calls. The datasets without benchmark VCF files will be used for speed evaluation. The following 5 metrics will be used for performance evaluation:

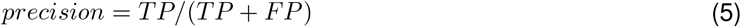

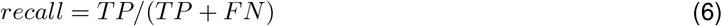

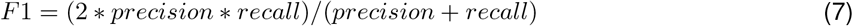

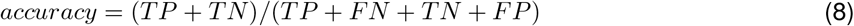

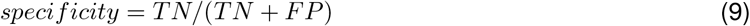

where TP refers to true positives, TN refers to true negatives, FP refers to false positives, and FN refers to false negatives. Elapsed time, in either minutes or seconds, will be used for comparing the speed of the models. From now on, “r_artifact_removal” and “py_artifact_removal” will be used to refer to the R and Python model respectively before applying deep learning while “DL_v11” will be used to refer to the Python model after applying deep learning. These terms are mainly used in the comparison of the model results pre and post-deep learning. The software, which outputs additional analyses and VCF files beyond the final trained model, is still named FINDEL.

The initial version of the DL_v11 model was trained on the HCC1954 cell line dataset. Although both performance and speed results are shown in Table II, the main focus is on the difference in speed between the models written in Python vs R programming language. The elapsed time of py_artifact_removal is less than 3 minutes while the elapsed time of r_artifact_removal is 305 minutes. This is a significant speedup of more than 100 times while maintaining similar performance. The objective function value of py_artifact_removal is 80.34 while the objective function value of r_artifact_removal is 80.67. The higher the value, the better the performance. The difference is negligible in this case. Meanwhile, DL_v11 achieves a significantly higher objective function value of 89.21 with a training and inference time of 20 and less than 3 minutes respectively. However, the performance improvement based on this dataset alone is not robust enough. As seen from the table, there is very little labeled data available. Only 214 variants are labeled with 213 of them being classified as positive SNPs and only 1 negative SNP. From a supervised deep-learning perspective, this is an imbalanced classification problem and the performance metrics might be biased. Even though DL_v11 correctly predicted 211 out of 213 positive SNPs, it failed to predict the only negative SNP. It can maximize the objective function value simply by predicting all SNPs to be positive as they are the majority of the labels. This poses a problem when we want to minimize false positives. On the other hand, py_artifact_removal can detect the only negative SNP but misses much more positive SNPs, achieving an overall lower objective function value. Therefore, the other labeled datasets are used to provide a more robust estimate of the model performances. The key takeaway here is the superior speed achieved by the models written in Python as compared to R. The subsequent evaluation comparison will be made between the 2 python models: py_artifact_removal and DL_v11.

The subsequent versions of the DL_v11 models were trained on the GIAB consortium reference samples (HG001-HG007). The following results were based on the first method of training using the same VCF file for both training and testing. The performance comparison between py_artifact_removal and DL_v11 based on the 5 metrics is given in Appendix A. DL_v11 performed better on all the metrics (F1, accuracy, precision, specificity) except recall.

The performance comparison within py_artifact_removal is given in Appendix B. Generally, higher scores were achieved on the human genomes from the Ashkenazi Jewish (HG002/HG003/HG004) ancestry than the Utah/European (HG001) and Han Chinese (HG005/HG006/HG007) ancestries for F1, accuracy, and precision. For recall, similar scores were achieved on the human genomes from all the ancestries. Results were mixed for specificity. The performance comparison within DL_v11 is given in Appendix C. Generally, higher scores were achieved on the human genomes from the Ashkenazi Jewish ancestry than the Utah/European and Han ancestries for F1, accuracy, and precision, and recall. Results were mixed for specificity.

The next set of results used the second training method where the train and test set consisted of different human genomes. The F1, accuracy, and precision performance comparisons within DL_v11 for different train sets are given in Appendix D, E, and F respectively. Generally, higher scores were achieved on the human genomes from the Ashkenazi Jewish ancestry than the Utah/European and Han Chinese ancestries regardless of the train set used. The recall performance comparison is given in Appendix G. Generally, higher scores were achieved on the human genomes from the Ashkenazi Jewish and Utah/European ancestries than the Han Chinese ancestry regardless of the train set used. The human genome from the Utah/European ancestry (HG001) had the best recall for all the train sets. The specificity performance comparison is given in Appendix H. Generally, higher scores were achieved on the human genomes from the Ashkenazi Jewish ancestry than the Han Chinese ancestry regardless of the train set used. The human genome from the Utah/European ancestry (HG001) had the worst specificity for all the train sets. The average performance comparison using different train sets is given in Appendix I. Generally, higher average scores on all 5 metrics were achieved when the human genomes from the Han Chinese ancestry were used as the train samples. The time comparison between the 3 models is given in Appendix J. All datasets used here are from the FFPE and fresh frozen WES samples. On both datasets, SRR2911438 and SRR2911453, a significantly shorter time was used by the 2 models written in Python (py_artifact_removal and DL_v11) as compared to the model written in R (r_artifact_removal). The correlation between elapsed time and the number of SNPs for py_artifact_removal is given in Appendix K. All datasets used here are from the FFPE and fresh frozen WES samples and the breast cancer WES samples. A strong positive linear correlation can be observed. As the number of SNPs increases, the elapsed time taken by py_artifact_removal increases as well.

## 4 Discussion

FINDEL, a variant refinement software infused with deep learning, has demonstrated its ability to efficiently remove sequencing artifacts from cancer samples using mutational signatures and other features. We have evaluated its performance and speed on high-quality open-source datasets. The performance is largely boosted by the supervised deep learning approach in addition to the use of mutational signatures. False positives are greatly reduced through the variant refinement process. The algorithm speed is mostly due to the optimal choice of programming language and code refactoring. The overall software runtime is low as our approach queries VCF files instead of BAM files where the former has a much lower memory footprint. Moreover, the software can be easily run on a local machine. FINDEL eliminates the need for users to manually inspect BAM files using some form of genomics viewer program. This saves both labor and time, increasing the turnover of cancer sample interpretation. Doctors require less time for the clinical diagnosis of patients and can increase the range of therapeutic opportunities for them. More importantly, the automation and standardization of the variant refinement process increase reproducibility by different entities. The usage of mutational signatures to partition the unrefined mutation set into refined and artifactual mutation subsets was possible as various artifacts had characteristic mutational signatures that were different from the true variants. These mutational signatures are constantly updated and can be found in the COSMIC database. This essentially removes the need for bioinformaticians to manually inspect large BAM files and search for patterns that are representative of artifacts using domain knowledge. The manual inspection process is a major bottleneck impeding the progress of the downstream analyses after the variant calling process. Eliminating this laborious task greatly reduces the entire bioinformatics workflow time. As the research on mutational signatures become more comprehensive in the future, we expect the algorithm to further improve in performance as well. FINDEL only requires a user-provided VCF file and an SBS mutational signature data file. No manual preprocessing on the user part is required as long as the standard format is used. Subsequently, FINDEL outputs a VCF file representing the refined mutations. The infusion of supervised deep learning to FINDEL mainly aims to improve algorithmic performance and further reduce the number of false positives. It also utilizes the information from the VCF files beyond just the mutational signatures. Specifically, additional features are engineered from the “INFO” and “SAMPLE” columns of the VCF files. As seen from the results, the addition of supervised deep learning greatly improves the score of most of the metrics. The disadvantage is that the initial training of the neural network might take quite some time. However, subsequent usage of the model, aka inference, takes little time. The deep learning model only needs to be retrained if there is evidence of model degradation. Also, the performance of the model is affected by the quality of the data. Generally, higher scores were achieved on the human genomes from the Ashkenazi Jewish ancestry than the Utah/European and Han Chinese ancestries due to the former having higher sequencing depth.

## 5 Conclusions and Future Work

We have developed a deep learning-based bioinformatics software, FINDEL, that can efficiently remove sequencing artifacts from cancer samples using mutational signatures and other features. It eliminates the laborious process of manually inspecting large BAM files by querying much smaller VCF files. The algorithm automates and standardizes the variant refinement process, reducing required labor and time while increasing reproducibility. The user only needs to provide the input files and the software does the rest of the work. Within minutes, the unrefined VCF file containing the artifactual variants are removed and the refined VCF file can be used for subsequent downstream analyses. The potential of FINDEL to redefine the bioinformatics workflow through its superior performance and speed allows better patient management and therapeutic opportunities. As of now, the software mainly focuses on single nucleotide polymorphisms (SNPs). In the future, the scope can be broadened to insertions and deletions, copy number variations, and even structural variations. Overall, we show that the use of mutational signatures coupled with deep learning can replace the manual inspection process, potentially setting a new standard for the variant refinement process.

## 6 Data availability

The datasets can be downloaded at their original sites.

## 7 Code availability

The source codes are only available under research or commercial collaboration.

## 8 Web service

We set up a web service hosting our trained model.

## 9 Acknowledgements

Not applicable.

## 10 Author contributions

